# Arabidopsis Membrane Bound O-acyl Transferase GHL1 is an essential orthologue of fungal GUP1 GPI-anchor remodelases and metazoan HHAT N-palmitoyl transferases

**DOI:** 10.64898/2026.08.27.744591

**Authors:** Yelizaveta Prokhorova, Sonica Chaudhry, Krzysztof Wypijewski, Sophie Cooke, Cairo Davidson, Mitzi Yoong, Jens Tilsner, Piers A. Hemsley

## Abstract

The GUP1/HHAT family of MBOAT proteins have been implicated in GPI-anchor acyl-chain remodelling in yeast and rice and secreted peptide acylation in eumetazoans, but whether these activities are truly distinct or GUP1/HHAT proteins are bifunctional has not been addressed. We show that the GUP1/HHAT family form a distinct clade within eukaryotic MBOATs with clear conservation of structural features, suggesting a single evolutionary event for their origin and a common mode of action. However, HHAT-like proteins form a clear and separate eumetazoan-unique group within the clade. This, in the context of information from other eukaryotes, suggests that GUP1-like GPI-anchor remodelase activity is the ancestral function, with eumetazoan HHAT-like peptide acyl-transferase activities emerging via gain-of-function. Genetic and cell biological examination of Arabidopsis GHL1 revealed that GHL1 localises to the Golgi and that Arabidopsis plants homozygous for loss of GHL1 cannot be recovered. Reciprocal cross analysis revealed severe effects on transmission, in particular through the male gamete. This is a common and well characterised phenotype of mutants in the Arabidopsis GPI-anchor synthesis pathway, supporting the notion that Arabidopsis GHL1 is a GPI-anchor remodelase rather than peptide acyl transferase. By reconstituting HHAT peptide acyl transferase activity towards generic Hedgehog-like peptides in plants we demonstrate that Arabidopsis GUP1 AND HHAT-LIKE 1 (GHL1) likely does not possess appreciable peptide acyltransferase activity and that peptide acyltransferase activity cannot easily be introduced via mutation of key residues to mimic the HHAT active site. Through this work we provide a potential new route to cell surface display of proteins in eukaryotic systems using co-expression of HHAT with the minimal acyl-acceptor peptide sequence of Hedgehog morphogens fused to the protein of interest.

## Introduction

Glycophosphatidylinositol (GPI)-anchored proteins are found on the extracellular leaflet of the plasma membrane and are involved in a diverse range of processes in plants, from regulating cell wall deposition and rearrangement to determinising self-incompatibility and pathogenesis (reviewed (Yeats et al., 2018)). Defects in GPI-anchor synthesis have major (Lalanne et al., 2004; Tashima et al., 2006) effects in Arabidopsis, humans and yeast, indicating their essential role in cellular function. In Arabidopsis, the importance of GPI anchors is particularly well characterised through their effects on cell wall production, fertility and embryogenesis, with many knock-out mutants in GPI anchor synthesis and remodelling enzymes being functionally lethal (Desnoyer & Palanivelu, 2020; Lalanne et al., 2004). GPI anchors are complex glycolipid moieties, initially synthesised on the cytosolic face of the endoplasmic reticulum (ER) before being flipped across the ER membrane onto the lumenal face. Within the ER lumen nascent GPI-anchors are extensively remodelled before being attached to soluble proteins within the ER lumen that contain a C-terminal ω-sequence (Gerber et al., 1992). This anchors the protein to the lumenal membrane leaflet, and the protein then proceeds from the ER through the secretory pathway via the Golgi to the cell surface. During this progression, remodelling of the acyl chains occurs. In animals this is restricted to de-acylation/re-acylation reactions to produce a saturated long chain (C14:0-18:0) di-acyl glycerol lipid group in the final GPI anchor, but in plants and fungi very-long chain fatty acids (VLCFA, typically C26:0) are found at the di-acyl glycerol *sn-2* position, and are added by the Membrane Bound O-Acyl Transferase (MBOAT) family Glycerol Uptake 1-like (GUP1-like) protein (Bosson et al., 2006; Xu et al., 2022).

Medium and long chain acylation of secreted peptides, such as Ghrelin, Wnt, Spitz and Hedgehog family members, has been described in many eumetazoans, with the identified enzymes all belonging to the MBOAT family of proteins (Buglino & Resh, 2008; Miura et al., 2006; Takada et al., 2006; Yang et al., 2008). In particular, Hedgehog acylation by palmitate (C16:0) is performed by Hedgehog acyltransferase (HHAT) (Buglino & Resh, 2008), a homologue of GUP1-like proteins, suggesting functional diversification or dual activity within this MBOAT family. An interesting report suggests that human and *Drosophila melanogaster* HHAT is able to partially complement the GPI-anchor related defects of *Candida albicans* mutants lacking GUP1 (Lucas et al., 2016), suggesting that as well as being a peptide acyltransferase HHAT may also be a GPI-anchor remodeller. This raises the intriguing counter question: do GUP1-like proteins, classified as GPI-remodellers, also have undocumented (secondary) peptide acyl transferase activity, raising the possibility of extracellular acylated polypeptides in plants and fungi?

## Results

### At1g57600 is a member of the HHAT/GUP1 MBOAT family

Membrane Bound O-Acyl Transferases have been identified in all groups of the eukaryotic phylogeny, typically adding lipids to substrates through an ester linkage. The MBOAT GUP1 from yeast and HHAT from eumetazoans have been proposed as homologues based on sequence (Lucas et al., 2016) despite differing in their published activities; GUP1 is a GPI anchor remodeller (Bosson et al., 2006) while HHAT acts to add fatty acyl groups to secreted Hedgehog peptides (Buglino & Resh, 2008). Using BLAST we identified At1g57600 as the closest reciprocal amino acid sequence match to both yeast GUP1 and human HHAT in Arabidopsis and renamed At1g57600 as GUP1 AND HHAT-LIKE 1 (GHL1). Amino acid-based phylogenetic analysis of all MBOATs from yeast, humans and Arabidopsis, plus validated GUP1 and HHAT-like activities from *Trypanosoma brucei* (Jaquenoud et al., 2008), *Candida albicans* (Ferreira et al., 2010), *Oryza sativa* (Xu et al., 2022) and *Drosophila melanogaster* (Buglino & Resh, 2008), indicates that GUP1-like and HHAT-like activities share a common ancestor. Arabidopsis GHL1 is placed firmly within the validated GUP1-like activity group that includes rice, fungi (yeast and *Candida albicans*) and *Trypanosoma brucei*, and GHL1 groups closest to the rice GPI remodelase BC16 (figure 1A). HHAT activities from humans and *Drosophila* form a clear sister group. This arrangement does not reflect the accepted view that Archaeplastida (including plants) and Discoba (e.g., *T. brucei*) diverged from the Opisthokont (fungi and animals) common ancestor long before the fungi-animal split (Gabaldon, 2021). This apparent contradiction indicates that HHAT-like activity likely arose through gain-of-function mutations in the ancestral GUP1-like activity in a eumetazoan progenitor organism.

**Figure 1.**
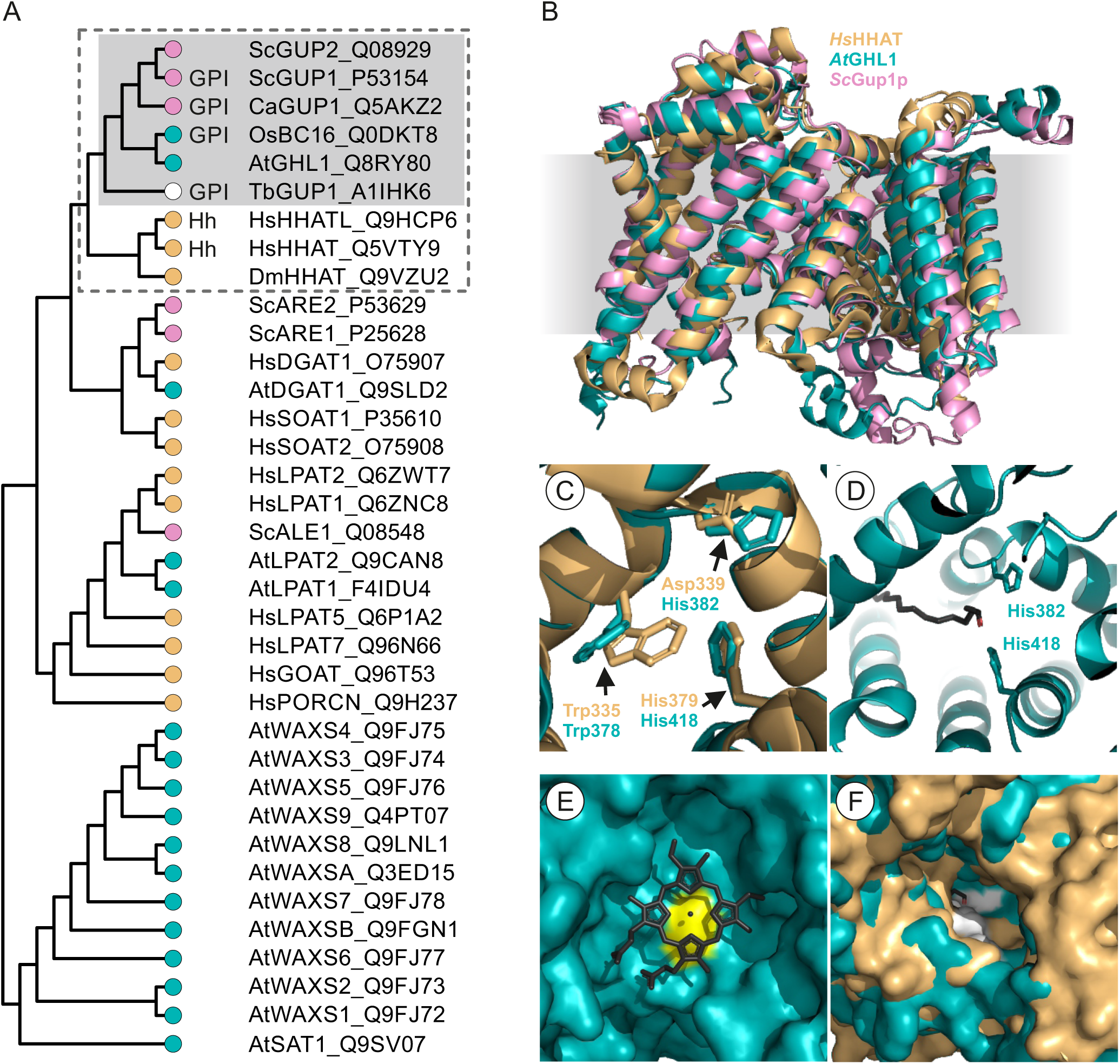
Arabidopsis GHL1, yeast GUP1 and human HHAT are homologues. **A.** Unrooted cladogram showing relations between plant (teal), fungal (pink), animal yellow and discobid (white) MBOAT family members. Validated activites are denoted by GPI for GPI anchor remodelase or Hh for peptide acyl transferase activity. The GUP1/HHAT-like clade is surrounded by a dotted line, with the sub-group containing validated GPI-remodelase activities boxed in grey. At - *Arabidopsis thaliana,* Ca – *Candida albicans,* Dm – *Drosophila melanogaster,* Hs – *Homo sapiens*, Os – *Oryza sativa,* Sc – *Saccharomyces cerevisiae,* Tb – *Trypanosoma brucei*. Uniprot accession numbers are shown after each protein name **B.** Structural comparison of Arabidopsis GHL1 and yeast GUP1 AlphaFold2 models with a cryo-EM determined structure of HHAT. **C.** Catalytically important residues are conserved between human HHAT (yellow) and Arabidopsis GHL1 (teal). **D.** Active site histidines in Arabidopsis GHL1 are positioned to coordinate fatty acids (black) within the acyl chain tunnel. **E.** The human HHAT Haem coordinating cysteine residue (bright yellow) and haem docking site are conserved in Arabidopsis GHL1. Haem shown in black. **F.** The luminal substrate access tunnel is conserved between human HHAT yellow and Arabidopsis GHL1 (teal). Active site residue shown in white, fatty acid coloured black.

Despite their divergent function, subsequent structural comparisons using the cryo-EM structures of HHAT (Coupland et al., 2021; Jiang et al., 2021) and alpha-fold predictions of Arabidopsis GHL1 and yeast GUP1 revealed a remarkable degree of structural conservation (figure 1B), including the positioning of the “gatekeeper” residue (Trp376 in GHL1, Trp335 in HHAT; figure 1C), acyl chain tunnel (figure 1D), haem-binding pocket and haem co-ordinating Cys-Arg dyad (Cys367/Arg293 in GHL1, Cys324/Arg250 in HHAT; figure 1E), lumenal substrate access pocket (figure 1F), and S-acylation sites, confirming that these proteins are likely closely related. However, in HHAT the active site residues consist of a unique Asp339/His379 dyad, while in GHL1 and all other GUP1-like proteins the equivalent amino acids are His/His (His382/His418 in GHL1; figure 1C) suggesting a different reaction mechanism. In HHAT, the Glu59 side chain carboxyl group forms a crucial ionic interaction with the conserved Arg28 side chain amino group of the Sonic Hedgehog peptide (Hardy & Resh, 2012; Jiang et al., 2021). In GHL1 the corresponding residue is Gln65 (neutral, amide side chain) suggesting that GHL1 is unlikely to use a similar substrate binding mode.

### GHL1 is essential for full male gamete viability

During our work, an investigation into the Rice homologue of GHL1, Brittle Culm 16 (BC16), was published. The genetic lesion in the *bc16* mutant introduces a stop codon after 225 amino acids (∼1/3 of the way through the coding sequence), is likely functionally null, and leads to dwarfing and fragile internodes (Xu et al., 2022). To compare phenotypes between rice and Arabidopsis we identified a number of T-DNA insertion lines in GHL1 (figure 2A). For most (SALK_015602, SALK_121939, SALK_015594 (Alonso et al., 2003), SAIL_518_E08 (Sessions et al., 2002), GABI_724B03 (Kleinboelting et al., 2012)) we were able to recover homozygotes for the insertion in GHL1, but were unable to observe any gross phenotype or effects on GHL1 expression; these lines were not studied further. However, for SALK_015690 and SALK_073335 we were never able to recover homozygous plants, despite repeated screening of offspring from heterozygous plants. Introduction of a *GHL1-3xMYC-EGFP* fusion expressed from the *35S* promoter complemented this defect and allowed for recovery of plants homozygous for the insertion in SALK_073335 in the T_3_ generation (figure 2B), indicating that the observed defects in homozygous mutant recovery are due loss of GHL1 function. SALK_015690 and SALK_073335 were therefore renamed *ghl1-1* and *ghl1-2* respectively. Kanamycin resistance-based screening through the *nptII* resistance gene carried on the inserted T-DNA in *ghl1-2* showed a Kan^Resistant^:Kan^Sensitive^ ratio of 372:497 (simplifies to ∼0.75:1, expect 3:1 for classical mendelian inheritance at a single locus). This indicates that there may be an effect, potentially involving gene dosage, on embryogenesis and/or transmission through one, other or both gametes. *ghl1-1* showed no resistance to kanamycin, likely due to *nptII* gene silencing often observed in SALK T-DNA lines (Alonso et al., 2003), precluding similar analysis. Introduction of the functional *GHL1-3xMYC-EGFP* fusion expressed from the embryo active *ABI3* promoter (Bodi et al., 2012) did not allow recovery of homozygotes. This suggests that the failure to recover homozygous *ghl1-2* seed is likely due to a defect in one or other gametophyte rather than (solely) embryo formation and development.

**Figure 2.**
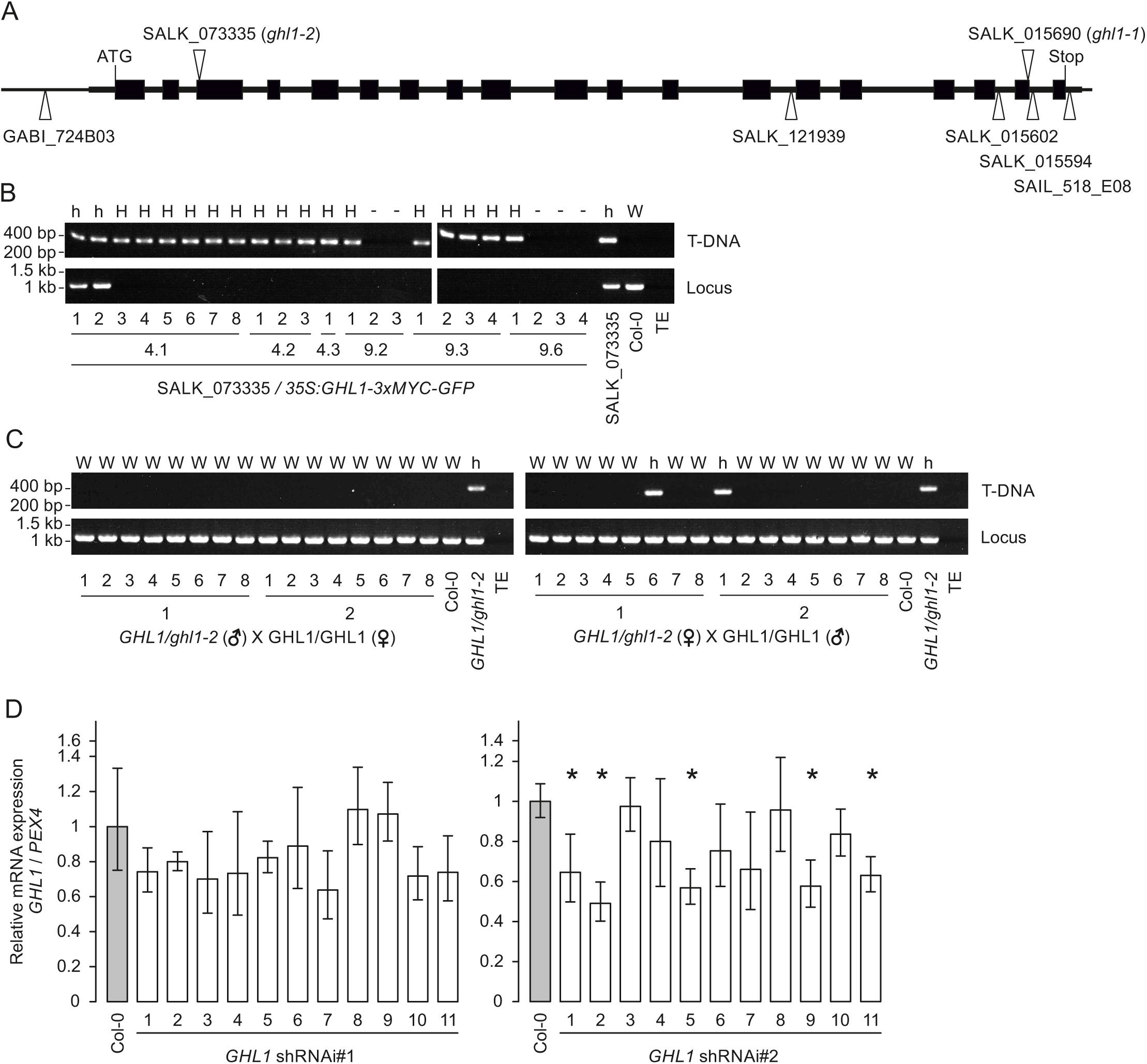
Loss of function mutants in GHL1 abolish transmission through the male gamete. **A.** Intron/exon structure of At1g57600 showing positions of T-DNA insertions. **B.** Transformation of plants heterozygous for the *ghl1-2* SALK_073335 T-DNA insertion with *35S:GHL1-3xMYC-GFP* allows for recovery of plants homozygous for the SALK_073335 T-DNA insertion. T_3_ plants from 2 independent transformants were analysed by PCR using primers specific for the T-DNA left border and the wild type At1g57600 genomic locus. h = heterozygous at locus, H= homozygous mutant at locus, W = wild type, - = no data obtained. **C.** *ghl1-2* shows no transmission through the male and reduced transmission through the female. DNA extracted from *GHL1/ghl1-2* x *GHL1/GHL1* (Col-0) F_1_ offspring was analysed by PCR using primers specific for the T-DNA left border and the wild type genomic locus. Data from 2 independant reciprocal crosses are shown. **D.** shRNAi does not allow recovery of plants with highly reduced *GHL1* transcript levels as determined by qRT-PCR. Data from T_1_ Col-0 plants transformed with two separate shRNAi constructs targeting the GHL coding sequence are shown. Values were calculated using the ΔΔCT method from 3 technical replicates, error bars represent RQMIN and RQMAX and constitute the acceptable error level for a 95% confidence interval according to Student’s t-test: * denotes plants with a significant reduction in *GHL1* transcript.

Reciprocal crosses between Col-0 and *ghl1-2* heterozygotes indicated that when *ghl1-2* heterozygotes were used as the female (figure 2B) ∼20% of offspring (7/35 individuals from 4 separate crosses) carried the T-DNA (expect 50% if no effect on female gamete) while no offspring carrying the T-DNA were observed when *ghl1-2* heterozygotes were used as the male (38 individuals tested from 6 separate crosses, expect 50% if no effect on male gamete, figure 2B). This suggests that while loss of *GHL1* affects formation or viability of female gametes to some degree, it is essential for male gamete development, growth or competitiveness; a classic phenotype of GPI anchor biosynthesis mutants in Arabidopsis (Desnoyer & Palanivelu, 2020) and further supporting a role for GHL1 in GPI anchor biosynthesis. As rice BC16 (Xu et al., 2022) and Arabidopsis GHL1 are single copy genes with no apparent paralogues, these data suggest that, in contrast to rice, loss of *GHL1* is highly detrimental to Arabidopsis and suggests a differential requirement for GPI anchored protein remodelling during gamete formation between plant species.

To try and observe whether reduction, rather than outright loss, of GHL1 would reveal informative somatic phenotypes we generated two pRNAi-GG (Yan et al., 2012) based small hairpin RNAi constructs targeting different parts of the *GHL1* sequence (RNAi#1 nucleotides 575-803, RNAi#2 nucleotides 1283-1480 relative to A of the start codon in *GHL1* cDNA) and transformed each into Col-0 plants. Pooled Kanamycin resistant T_2_ seedlings from each line were tested for expression of *GHL1.* Many *GHL1 RNAi#1* plants showed reduced but non-significant reduction in transcript levels, while half of all lines from *GHL1 RNAi#2* plants showed a reduction to approximately 50% of Col-0 levels (figure 3E). However, none of these lines showed any phenotypes at any life stage examined, suggesting that the partial knockdown achieved does not reduce GHL1 activity to a level that impacts cellular function, similar to heterozygous *ghl1-1* and *ghl1-2* which also display no observable somatic phenotype. It is plausible that plants with stronger knock down effects may have been generated but would, to all intents and purposes, be sterile due to deleterious effects on gamete viability and are therefore not recovered.

**Figure 3.**
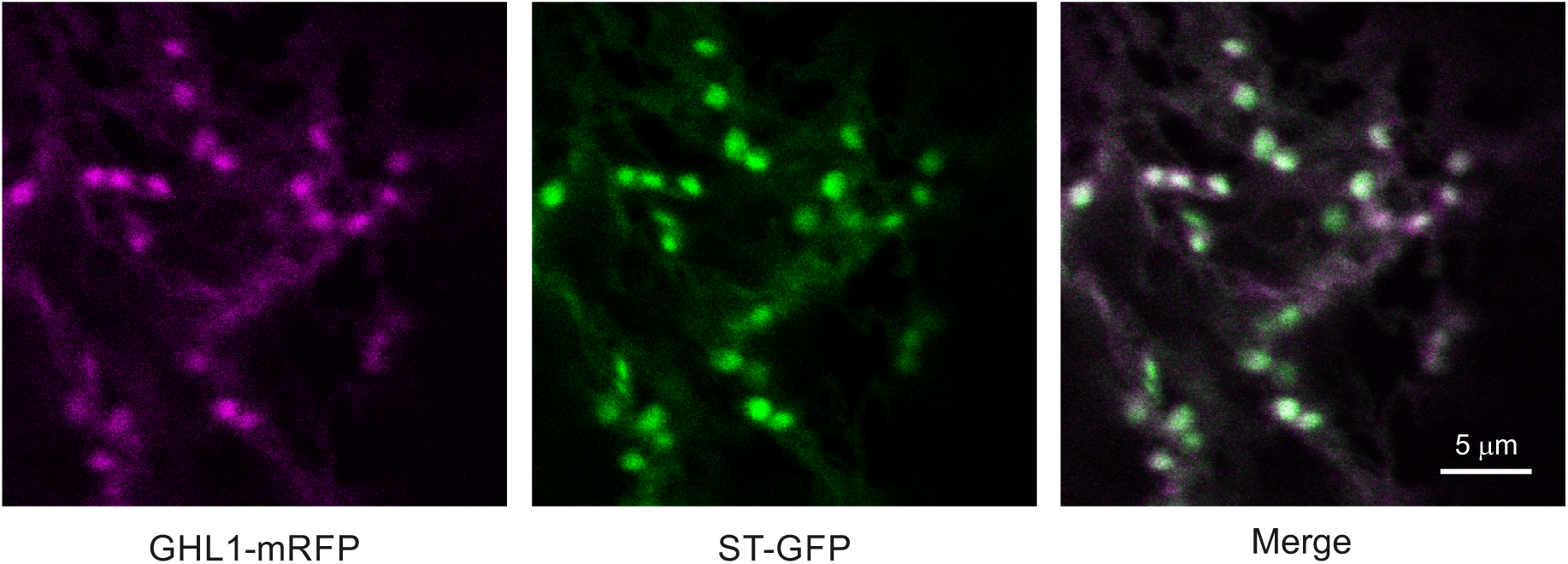
GHL1 primarily localises to the Golgi. GHL1-mRFP fusions show a high degree of overlap with a Sialyl transferase-GFP Golgi marker when co-expressed in *N. benthamiana*. Image shows a single slice from a z-stack. Green = GFP channel, Magenta = mRFP channel.

### GHL1 localises to the Golgi

If, as seems likely based on homology and phylogeny, GHL1 is involved in GPI-anchor remodelling it would be expected to reside in the secretory pathway. To test this, we co-infiltrated GHL1-mRFP and ST-GFP (Sialyl Transferase-GFP; Golgi marker (Boevink et al., 1998)) and observed a weak reticulate distribution, typical of the ER, in the mRFP channel, with bright foci showing dual mRFP/GFP labelling, indicating a predominant Golgi localisation of GHL1 (figure 3) identical to that reported for Rice BC16 (Xu et al., 2022).

### HHAT but not GHL1 demonstrates peptide acyl transferase activity in planta

Despite their remarkable structural conservation, within the GUP1/HHAT-like family there appears to be a functional divergence between those that perform GPI-anchor remodelling, best characterised using yeast GUP1 (Bosson et al., 2006) and recently demonstrated for rice BC16 (Xu et al., 2022), and peptide acyl-transferase activity in eumetazoans, originally identified in Drosophila HHAT (Buglino & Resh, 2008). Phylogenetic analysis (Figure 1A) suggests that HHAT peptide acyl-transferase activity is likely a derived function, as evidence for GUP1 GPI remodelling activity is found in rice (Xu et al., 2022), trypanosomes (Jaquenoud et al., 2008) and fungi (Bosson et al., 2006) but HHAT-like peptide acyl transferase activity has only been reported in eumetazoans. Interestingly, loss of Hedgehog substrate in Drosophila is milder than, and does not recapitulate, the observed phenotype of mutants in HHAT, suggesting that HHAT has additional (e.g. SPITZ (Miura et al., 2006)) or as yet unidentified substrates, or is bifunctional as a peptide acyl transferase and GPI-remodelase. In support of this latter idea both mouse and Drosphila HHAT are able to partially complement GPI-anchor related defects of *Candida albicans* GUP1 mutants (Lucas et al., 2016). This raises the question of whether conversion of a GUP1-like GPI-remodelase activity into a HHAT peptide acyl-tranferase-like activity is a simple process.

To determine whether HHAT activity can be reconstituted in plants, we expressed *Drosophila melanogaster* HHAT (*Dm*HHAT) alongside a minimal synthetic Drosophila Hedgehog substrate (Buglino & Resh, 2008) (minHh: <u>C</u>GPGRGLGRG; acyl acceptor Cys underlined) fused to a C-terminal 1xFLAG-GS-mRFP sequence in *N. benthamiana*. Using Triton X-114 phase partitioning (Bordier, 1981) as a readout for Hedgehog acylation (Miura et al., 2006) we found that minHh-1xFLAG-GS-mRFP was almost exclusively found in the detergent phase in the presence of *Dm*HHAT but not when expressed on its own (Fig 4A). This indicates that minHh acylation had occurred and that HHAT is catalytically active towards its native Hedgehog substrate in plants without any additional non-plant cofactors being required. A second mRFP antibody reacting band was observed almost exclusively in the soluble fraction at the expected molecular mass for mRFP. This band was not detected using anti-FLAG antibodies suggesting that proteolytic cleavage had occurred between the mRFP epitope and minHh sequence, releasing free, soluble mRFP without a FLAG tag. To test whether GHL1 had acyl transferase activity towards Hedgehog we co-expressed GHL1 with minHh but observed no minHh signal in the detergent phase, indicating that acylation had not occurred (Fig 4B). Two features of HHAT proposed to be necessary for effective Hedgehog acylation, the catalytic Asp/His dyad and the Glu59 Hedgehog Arg28 engaging residue (Gln in GHL1) (Coupland et al., 2021), are not conserved in GHL1. We therefore designed mGHL1 containing H^382^D and Q^65^E mutations to mimic these critical HHAT residues, and potentially recapitulate the evolutionary steps required to convert GUP1-like GPI remodelase activity into HHAT-like peptide acyl transferase activity. However, we observed no mGHL1 acylation activity towards minHh (Fig 4B). This may be due to co-evolution of HHAT and Hedgehog sequence/structure to interface effectively and promote acylation. However, the absolute peptide sequence specificity of Drosophila HHAT for its minimal Hedgehog CGPGRGLGRG substrate is apparently low, with an N-terminal cysteine after signal peptide cleavage being the only absolute requirement for acylation, so long as the subsequent ∼5 amino acids can fit into the active site pocket, and a positively charged residue is present in the 4-6 amino acids after the cysteine to engage with HHAT Glu59 (Hardy & Resh, 2012). We theorised that the proline in the <u>C</u>GPGRGLGRG sequence may limit peptide flexibility/conformation and hinder potential GHL1/mGHL1 acylation activity towards the model substrate. We therefore tested minHh variants where the proline was mutated to glycine (minHh-P3G) or alanine (minHh-P3A) to introduce greater flexibility and limit steric constraints of the peptide but, again, observed no acylation by GHL1/mGHL1, while HHAT was fully effective against these artificial substrates (Fig 4C, D). We conclude that while HHAT activity against a range of minimal hedgehog substrates can be fully reconstituted *in planta*, GHL1 or mGHL1 is not able to act as a peptide acyl transferase within this experimental set up. Attempts to model Hedgehog-derived peptides within the HHAT, GHL1 or mGHL1 substrate cavity using AlphaFold2-multimer (Evans et al., 2021) through ColabFold (Mirdita et al., 2022) were inconclusive. In all cases the peptide was located in the approximate position and expected orientation for active site engagement within the GHL1/mGHL1/HHAT structure (pTM >0.8) but ipTM values were <0.7 for all GHL1/mGHL1 combinations with minHh variants and for the biologically relevant HHAT/minHh combination with an empirically determined structure (Coupland et al., 2021; Jiang et al., 2021) indicating a high degree of uncertainty when predicting absolute conformations at the site of interaction. However, overall MBOAT structure did not need to change to accommodate the substrate peptides (e.g., apo-GHL1 vs GHL1 with minHh: RMSD = 0.558 Å across all residues of GHL1) indicating that neither physical obstruction nor substrate accessibility likely explains the lack of GHL1/mGHL1 activity towards the peptides. While Hedgehog-derived peptides were placed in the correct orientation and general position for active site engagement to be possible in HHAT, GHL1 and mGHL1, confidence in overall minHh variant backbone and side chain positioning (pLDDT <60 for all minHh variants, including for the biologically relevant HHAT/minHh combination) prevented meaningful interpretation of predicted structures to provide mechanistic solutions to introduce Hedgehog-like acylating activity to GHL1.

**Figure 4.**
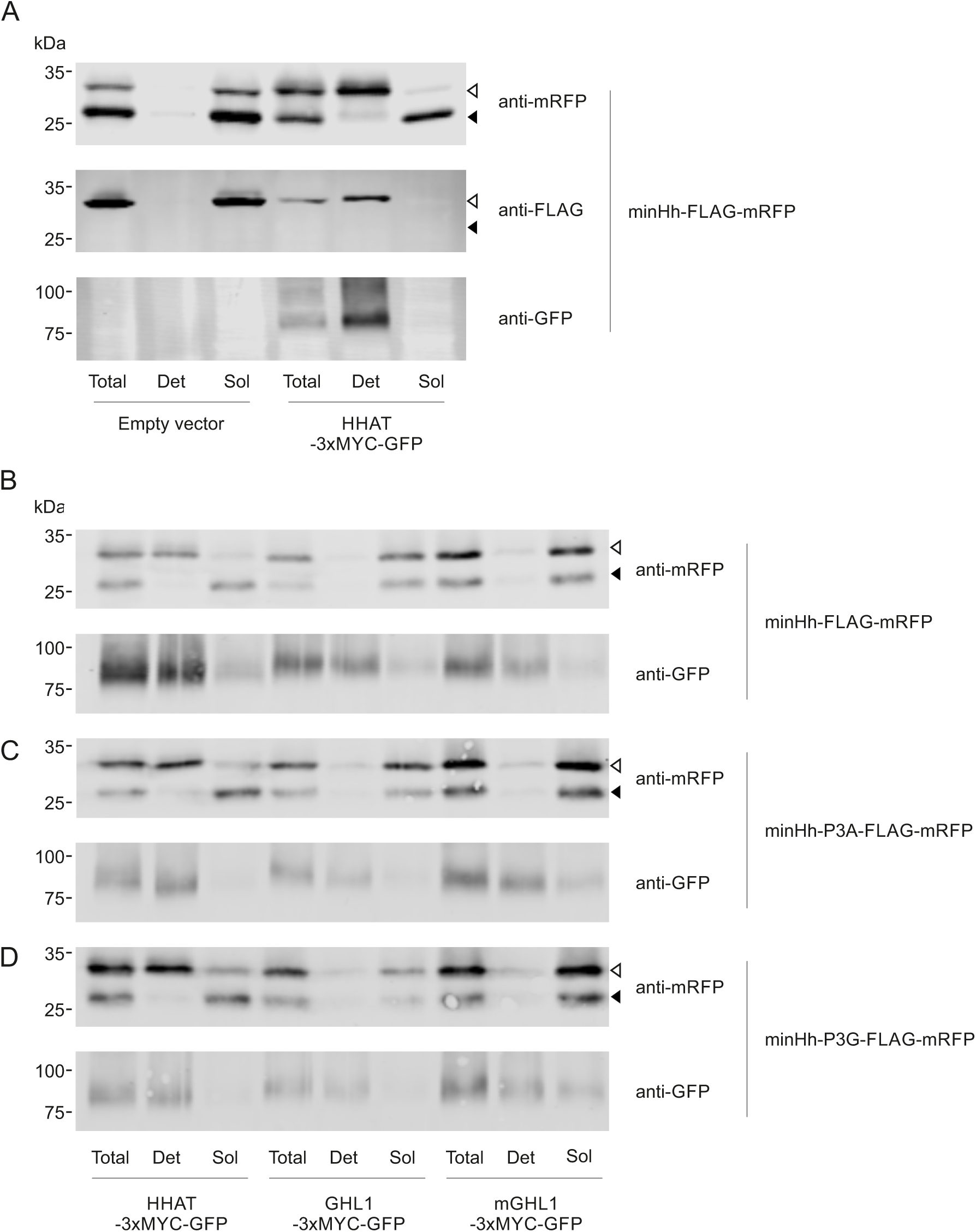
HHAT, but not GHL1, has peptide acyl transferase activity *in planta*. **A.** HHAT activity can be reconstituted in *N. benthamiana* and detergent phase enrichment of Hedgehog (minHh - CGPGRGLGRG fused N-terminal to FLAG-mRFP) requires HHAT. HHAT, but not GHL1 or mGHL1, is able to promote the acylation of **B.** Hedgehog (minHh - CGPGRGLGRG), **C.** Hedgehog-derived minHh-P3A - CGAGRGLGRG or **D.** minHh-P3G - CGGGRGLGRG peptides fused N-terminal to a FLAG-mRFP tag when transiently expressed in *N. benthamiana*. minHh variant acylation is assessed by Triton X-114 phase partitioning into the detergent (Det, acylated) or soluble (Sol, non-acylated) phases. Pre-fractionation total lysate is shown (Total) as an input reference. Upper band (open arrowhead) in mRFP blots is minHh_variant-FLAG-mRFP, lower band (shaded arrowhead) is free mRFP following cleavage within the FLAG-tag sequence and acts as a control for soluble phase enrichment. Presence of integral membrane HHAT/GHL1/mGHL1-3xMYC-GFP fusions in samples and phase separated fractions is shown by anti-GFP blot and acts as a control for detergent phase enrichment.

## Discussion

During the course of our work on GHL1 it emerged that rice BC16 encodes a GUP1-like protein with GPI anchor remodelase activity whose loss causes dwarfing and a range of cell wall defects likely due to disruption of GPI-anchor biogenesis (Xu et al., 2022). However, while mutation of GHL1 was apparently highly detrimental to male gametophyte formation, viability or growth, resulting in an inability to recover homozygotes, Rice BC16 mutants only showed a ∼25% reduction in seed set (Xu et al., 2022), indicating a less stringent requirement for GUP1-like activity in rice compared to Arabidopsis. It is possible that Arabidopsis GPI anchor proteins play a much greater role in male gametophyte maturation or growth than in rice; many other Arabidopsis GPI synthesis mutants are male sterile (Desnoyer & Palanivelu, 2020; Lalanne et al., 2004). Similar to rice BC16, we observed GHL1 in the Golgi and ER, lending support to their proposed function as GPI-remodellers in the secretory pathway.

Based on prediction and proteomics (Eisenhaber et al., 2003; Takahashi et al., 2016), approximately 150-250 proteins from Arabidopsis are thought to be modified by GPI anchors, and the generation and remodelling of these anchors is a multi-step process. After synthesis and sugar group remodelling, GPI-anchors are attached to proteins at defined C-terminal motifs as triply acylated precursors. POST-GPI ATTACHMENT TO PROTEINS 1 (PGAP1)/ BYPASS OF SEC THIRTEEN 1 (BST1) then removes the acyl group attached to the inositol ring (Bernat-Silvestre et al., 2021) and POST GPI ATTACHMENT TO PROTEINS 3 (PGAP3)/ PROCESSING IN THE ER (PER1) removes the acyl group from the *sn-2* position of diacylglycerol (Bernat-Silvestre et al., 2022) to generate *lyso*-phosphatidylinositol. In eumetazoans, reacylation of *lyso*-phosphatidylinositol is performed by POST GPI ATTACHMENT TO PROTEINS 2 (PGAP2) (Tashima et al., 2006), while in fungi and plants (Bosson et al., 2006; Xu et al., 2022) it appears to be performed by GUP1 homologues (e.g., yeast GUP1, rice BC16, likely Arabidopsis GHL1). Based on sequence and structure analysis, eumetazoan PGAP2 and plant or fungal GUP1 are not related and are functional analogues rather than homologues. In eumetazoans, the GUP1 homologue HHAT no longer appears to be essential for GPI anchor remodelling and instead adds acyl groups to secreted Hedgehog family peptides as its proposed primary function (Buglino & Resh, 2008; Miura et al., 2006). However, Drosophila and mouse HHAT are able to partially complement GPI-anchor related defects of GUP1 mutants in *Candida albicans* (Lucas et al., 2016), suggesting that some residual GPI anchor remodelling activity remains.

Structurally, GHL1, GUP1 and HHAT are clearly related (figure 1) and catalyse acyl-CoA mediated fatty acylations, yet they have differing acyl-acceptor substrates and bond chemistry. In the active site of HHAT the His/His dyad found in plant and fungal GUP1-like proteins is Asp/His. HHAT is unique amongst MBOATS in that the proposed acylation reaction proceeds via nucleophilic attack by the N-terminal Hedgehog cysteine amino group on the thioester of acyl-CoA, resulting in Hedgehog with an amide linked fatty acid (Jiang et al., 2021; Schonbrun & Resh, 2022), rather than the ester linked fatty acids typically found on other MBOAT reaction products. Asp339 is proposed to act as a general base to activate the N-terminal Hedgehog cysteine amino group (Jiang et al., 2021) but, based on our work, a His to Asp conversion at this site alone is not sufficient to impart detectable peptide acyl transferase activity to GHL1. Despite these differences, the close structural homology between GUP1-like proteins and HHAT should mean that mechanistic insights into GUP1, GHL1 and BC16 function can be made using information from HHAT structures.

In terms of how HHAT/GUP1-like function diverged, eumetazoan GPI anchors have not been reported to contain ceramides, unlike yeast and plants (Oxley & Bacic, 1999; Svetek et al., 1999). It has been noted previously that yeast CWH43, required for ceramide remodelling of GPI-anchors and reliant upon prior GUP1 mediated reacylation of *lyso*-phosphatidylinositol GPI anchors, shows homology at its N-terminus to eumetazoan PGAP2 (functionally analogous to GUP1 in terms of reacylating GPI anchors), and at its C-terminus to PGAP2 Interacting Protein (PG2IP) (Umemura et al., 2007). If ceramide containing GPI anchors provided no functional benefit to eumetazoans over saturated fatty acid GPI anchors, the relaxation in function of PGAP2 and/or PG2IP to mirror GUP1-like function in eumetazoans to transfer fatty acids as well as, or instead of, ceramides may have allowed the subsequent evolution of the ancestral eumetazoan GUP1-like protein into peptide acylating HHAT. Alternatively, a gain-of-function change of ancestral HHAT into a dual function GPI-remodeler and proto-peptide acyl-transferase may have placed selective pressure upon the eumetazoan progenitor organism to evolve alternative fatty acid transferase activity towards *lyso*-phosphatidylinositol; a role that filled by PGAP2/PG2IP, likely at the expense of ceramide transfer.

From a biotechnological perspective, heterologous reconstitution of HHAT activity, able to acylate a short peptide sequence derived from Hedgehog, offers a potential alternative route to cell surface display. Cell surface display is a powerful approach for immobilising proteins/peptides at the cell surface and facilitates, for example, nanobody library or biocatalyst screening, or enables bioactive peptide discovery, bioremediation, biosensor construction or therapeutic delivery (Li et al., 2024). Anchoring to the plasma membrane can also help to retain produced protein on the cell surface rather than having them diffuse into the media (cell culture) or apoplast (intact plants). Traditional approaches use a ∼25 amino acid C-terminal sequence for attachment of GPI anchors (Van der Vaart et al., 1997) or a ∼35 amino acid N-terminal tail anchor integral membrane peptide (Sueda et al., 2025). Here we demonstrate that the minimal 6-11 amino acid Hedgehog sequence could potentially also be used to anchor heterologous proteins to the outer leaflet of the plasma membrane so long as a HHAT-like activity is present in the cell (native or heterologous), and expands the options available for cell surface display in eukaryotic systems by providing a small, N-terminal lipid-based anchor, and will be particularly useful for proteins where a free C-terminus is required or where phospholipase C mediated cleavage of GPI anchors may be an issue.

## Methods

### Arabidopsis growth and transformation

All *Arabidopsis thaliana* work used Col-0 ecotype as wild type. T-DNA insertion lines in GHL1 (SALK_015690 (renamed *ghl1-1*), SALK_073335 (renamed *ghl1-2*), SALK_015602, SALK_121939, SALK_015594 (Alonso et al., 2003), SAIL_518_E08 (Sessions et al., 2002), GABI_724B03 (Kleinboelting et al., 2012) were obtained from NASC. Transgenic Arabidopsis plant lines were generated by *Agrobacterium tumefaciens* GV3101 pMP90-mediated floral dip transformation (Clough & Bent, 1998). Arabidopsis material was grown on 0.5x Murashige & Skoog (MS) medium containing 0.8% (w/v) plant agar. Seeds were stratified for 48h at 4°C and germinated under a 16/8 h light/dark cycle at 20°C in MLR-350 growth chambers (Panasonic). Phosphinothricin resistant Arabidopsis plants were initially selected on soil by spraying 10 days after germination, and subsequently at 2-day intervals, with Finale BASTA herbicide (120 µg/ml final concentration). Kanamycin resistant Arabidopsis plants, and BASTA resistant Arabidopsis plants from the T1 generation onwards, were selected on 0.5x MS containing 0.8% (w/v) plant agar supplemented with antibiotics (Kanamycin 50 µg/ml, Phosphinothricin 25 µg/ml, final concentrations) (Harrison et al., 2006).

### *Nicotiana benthamiana* growth and transient transformation

*Nicotiana benthamiana* plants were grown in commercial compost under glasshouse conditions of 16/8 h light/dark cycle, daytime 26°C, night-time 22°C for 4-5 weeks before use. *Nicotiana benthamiana* leaves were syringe infiltrated with *A. tumefaciens* GV3101 pMP90 containing expression plasmids for genes of interest alongside *A. tumefaciens* GV3101 pMP90 containing p19 (Voinnet et al., 2003), each at a final OD600 of 0.05-03 (in 10 mM 2-(N-morpholino)ethanesulfonic acid, 10 mM MgCl2, 150 µM acetosyringone, pH 5.7 in water) depending on observed expression levels in pilot experiments. Plants were maintained under standard growth conditions for a further 60h before analysis.

### Molecular biology

*GHL1* cDNA without stop codon (synthesised by Genscript) was used as PCR template for NEBuilder (NEB) assembly, either being cloned into *pENTR D-TOPO* to generate *pENTR GHL1*, or replacing the FLS2 coding sequence *in pENTR FLS2-3xMYC-GFP* (Hurst et al., 2023) to generate *pENTR GHL1-3xMYC-GFP*. *pENTR GHL1* was gateway LR recombined (Invitrogen) with *pGWB554* (Nakagawa et al., 2007) to generate *35S_pro_:GHL1-mRFP*. *pENTR GHL1-3xMYC-GFP* was gateway LR recombined with pB7WG0 (Karimi et al., 2002) to generate *35S_pro_:GHL1-3xMYC-GFP* and with *pGHABI3GWG* (Bodi et al., 2012) to generate *ABI3_pro_:GHL1-3xMYC-GFP*. *Drosophila melanogaster HHAT* was amplified from cDNA (Brown & Kafatos, 1988) by PCR and used with NEBuilder HiFi (NEB #E2621)) to replace the FLS2 coding sequence in *pENTR FLS2-3xMYC-GFP* (Hurst et al., 2023) to generate *pENTR HHAT-3xMYC-GFP*. *pENTR HHAT-3xMYC-GFP* was gateway LR recombined (Invitrogen) with pB7WG0 (Karimi et al., 2002) to generate *35S_pro_:HHAT-3xMYC-GFP.* Synthetic, minimal “hedgehog-like” substrates were generated by overlapping PCR using synthetic oligonucleotides and inserted into pENTR D-TOPO using NEBuilder HiFi before gateway LR recombination into *pGWB554* to create C-terminal mRFP fusions. GHL1 RNAi constructs were generated by PCR amplification of nucleotides 575-803 (#1) or 1283-1480 (#2) using *Bsa*I site containing GoldenGate adaptors before *Bsa*I GoldenGate assembly into pRNAi-GG (Yan et al., 2012).

### Gene expression analysis

Gene expression analysis was carried out using qRT-PCR as described [18, 36]. RNA was extracted from pooled 10-day old seedlings using an RNeasy mini kit with on-column DNAse digestion according to manufacturer’s instructions (QIAgen). cDNA was generated using 2 µg RNA with a high-capacity cDNA reverse transcription kit and random hexamer priming (Applied Biosystems). Expression levels were normalised against PEX4 (At5G25760, (Hemsley et al., 2014)). Relative quantification was calculated using the ΔΔCT method (Schmittgen & Livak, 2008). Error bars represent RQMAX and RQMIN over three technical replicates and constitute a 95% confidence interval (Student’s t-test).

### Triton X-114 phase partitioning

Based on the work described (Bordier, 1981; Miura et al., 2006), 6x leaf disc punches (#3 cork borer) from *N. benthamiana* plants expressing relevant constructs were lysed in 500 µl ice cold precleared 2% Triton X-114 (final, v/v) containing buffer (PBS, 5mM EDTA, protease inhibitors Merck P9599) using a micro pestle and solubilised at 4 °C for 25 minutes with gentle mixing. Samples were clarified by centrifugation at 16 kxg for 5 minutes at 4°C, supernatants retained and recentrifuged at 16 kxg for 5 minutes at 4°C. Protein concentration of supernatants was determined using BCA assay and samples diluted to 3 mg.ml^-1^ in ice cold lysis buffer. 1.5 mg of each sample was transferred to a 1.5 ml microfuge tube and placed in a 37 °C water bath for 30 minutes to allow phase separation to occur. Samples were fully phase separated by centrifugation at 16 kxg for 10 minutes at 24 °C. The clear, upper aqueous phase was removed to a fresh tube and the proteins in the aqueous and detergent phases were precipitated using 4 volumes of ice-cold acetone. Protein pellets were resuspended in 2x reducing Laemmli SDS-PAGE loading buffer before separation by SDS-PAGE and western blotting.

### Western blotting

Proteins were transferred to PVDF membrane using wet transfer and detected using monoclonal primary antibodies (α-GFP Roche 11867423001, α-FLAG M2 antibody Merck F1804 and α-mRFP ChromoTek RFP Monoclonal antibody 5F8) and Alexa Fluor® 790 AffiniPure Goat Anti-Mouse IgG (H+L) secondary antibody (Jackson ImmunoResearch 115-655-146). Signal was acquired and visualised using a Licor Odyssey CLx running Licor Image Studio Lite v5.2.

### Sequence and structural homology analysis

Amino acid sequences were obtained from Uniprot. Multiple alignment were performed using Clustal Omega (Sievers et al., 2011) and the resulting cladogram visualised using Iroki (Moore et al., 2020). Structures were analysed, aligned and compared using Pymol v2.5.5. Structure accessions: HHAT (PDB 7MHZ), GUP1 (AlphaFold AF-P53154-F1-model_v4), GHL1 (AlphaFold AF-Q8RY80-F1-model_v4).

### Microscopy

Transiently transformed *N. benthamiana* leaves were detached and immobilised on glass slides with double sided sticky tape, lower epidermis facing up. Imaging was performed on an upright Zeiss 710Meta confocal laser scanning microscope (Carl Zeiss, Jena) using a W Plan-Apochromat 40x/1.0 DIC M27 water dipping lens immersed in a drop of water placed directly on the plant sample. YFP was excited at 488 nm and mRFP at 561 nm. Detection ranges were 499 – 530 nm for YFP and 591 – 630 nm for mRFP, respectively. YFP and mRFP channels were imaged sequentially to minimize bleed-through. Images were acquired using ZEN software (Carl Zeiss, Jena) and exported as .TIF files.

Settings for figure 3 panels A-C were: 2% laser power for both excitation wavelengths; image size 8-bit 1024 x 1024 pixels at 3.5 x zoom corresponding to 60.73 x 60.73 µm; pixel dwell time 0.79 µs; pinhole 43 µm for both channels. YFP signal was pseudo-coloured green and mRFP signal magenta, both with linear LUTs covering the full range of data. The image was cropped with no further adjustments to resolution, brightness or contrast.

## Data availability statement

All data generated or analysed during this study are included in this published article or will be made freely available upon request. All physical research materials generated in this study will be made freely available upon request and completion of an MTA.

## Acknowledgements

We would like to thank Cândida Lucas (University of Minho) for critical discussions and advice during the preparation of this manuscript. We thank Jens Januske (University of Dundee) for providing the *Drosophila melanogaster* cDNA library used for amplification of *HHAT*. Joshua Booth performed work presented in this manuscript (see CRediT statement) but could not be contacted to approve authorship prior to submission. We would also like to thank Alasdair Munro and Sarah Dobbin for provision and/or maintenance of plant material used in this work. This work was supported by BBSRC grant BBSRC BB/W000261/1 and BB/R008787/1 to PH and Scottish Government Rural and Environment Science and Analytical Services (p) division funding to JT.

## CRediT statement

**Yelizaveta Prokhorova:** Validation, Formal analysis, Investigation, Data curation, Writing - Review & Editing. **Sonica Choudry**: Validation, Formal analysis, Investigation, Data curation, Writing - Review & Editing. **Krzysztof Wypijewski**: Validation, Formal analysis, Investigation, Data curation, Writing - Review & Editing. **Joshua Booth:** Investigation, Data curation, Formal analysis Writing - Review & Editing. **Sophie Cooke:** Investigation, Data curation, Formal analysis Writing - Review & Editing. **Mitzi Yoong**: Investigation, Formal analysis Data curation, Writing - Review & Editing. **Cairo Davidson**: Investigation, Formal analysis Data curation, Writing - Review & Editing. **Jens Tilsner**: Methodology, Investigation, Formal analysis, Data curation, Writing - Review & Editing, Visualization, Funding acquisition. **Piers A. Hemsley**: Conceptualization, Methodology, Validation, Formal analysis, Investigation, Data curation, Resources, Writing - Original Draft, Writing - Review & Editing, Visualization, Supervision, Project administration, Funding acquisition.

## References

Alonso, J. M., Stepanova, A. N., Leisse, T. J., Kim, C. J., Chen, H., Shinn, P., Stevenson, D. K., Zimmerman, J., Barajas, P., Cheuk, R., Gadrinab, C., Heller, C., Jeske, A., Koesema, E., Meyers, C. C., Parker, H., Prednis, L., Ansari, Y., Choy, N.,…Ecker, J. R. (2003). Genome-wide insertional mutagenesis of Arabidopsis thaliana. Science, 301(5633), 653–657.

Bernat-Silvestre, C., Ma, Y., Johnson, K., Ferrando, A., Aniento, F., & Marcote, M. J. (2022). Characterization of Arabidopsis Post-Glycosylphosphatidylinositol Attachment to Proteins Phospholipase 3 Like Genes. Front Plant Sci, 13, 817915. 10.3389/fpls.2022.817915

Bernat-Silvestre, C., Sanchez-Simarro, J., Ma, Y., Montero-Pau, J., Johnson, K., Aniento, F., & Marcote, M. J. (2021). AtPGAP1 functions as a GPI inositol-deacylase required for efficient transport of GPI-anchored proteins. Plant Physiol, 187(4), 2156–2173. 10.1093/plphys/kiab384

Bodi, Z., Zhong, S., Mehra, S., Song, J., Graham, N., Li, H., May, S., & Fray, R. G. (2012). Adenosine Methylation in Arabidopsis mRNA is Associated with the 3’ End and Reduced Levels Cause Developmental Defects. Front Plant Sci, 3, 48. 10.3389/fpls.2012.00048

Boevink, P., Oparka, K., Santa Cruz, S., Martin, B., Betteridge, A., & Hawes, C. (1998). Stacks on tracks: the plant Golgi apparatus traffics on an actin/ER network. Plant J, 15(3), 441–447. 10.1046/j.1365-313x.1998.00208.x

Bordier, C. (1981). Phase separation of integral membrane proteins in Triton X-114 solution. J Biol Chem, 256(4), 1604–1607.

Bosson, R., Jaquenoud, M., & Conzelmann, A. (2006). GUP1 of Saccharomyces cerevisiae encodes an O-acyltransferase involved in remodeling of the GPI anchor. Mol Biol Cell, 17(6), 2636–2645. 10.1091/mbc.e06-02-0104

Brown, N. H., & Kafatos, F. C. (1988). Functional cDNA libraries from Drosophila embryos. J Mol Biol, 203(2), 425–437. 10.1016/0022-2836(88)90010-1

Buglino, J. A., & Resh, M. D. (2008). Hhat is a palmitoylacyltransferase with specificity for N-palmitoylation of Sonic Hedgehog. J Biol Chem, 283(32), 22076–22088. 10.1074/jbc.M803901200

Clough, S. J., & Bent, A. F. (1998). Floral dip: a simplified method for Agrobacterium-mediated transformation of Arabidopsis thaliana. Plant J, 16(6), 735–743.

Coupland, C. E., Andrei, S. A., Ansell, T. B., Carrique, L., Kumar, P., Sefer, L., Schwab, R. A., Byrne, E. F. X., Pardon, E., Steyaert, J., Magee, A. I., Lanyon-Hogg, T., Sansom, M. S. P., Tate, E. W., & Siebold, C. (2021). Structure, mechanism, and inhibition of Hedgehog acyltransferase. Mol Cell, 81(24), 5025–5038 e5010. 10.1016/j.molcel.2021.11.018

Desnoyer, N., & Palanivelu, R. (2020). Bridging the GAPs in plant reproduction: a comparison of plant and animal GPI-anchored proteins. Plant Reprod, 33(3-4), 129–142. 10.1007/s00497-020-00395-9

Eisenhaber, B., Wildpaner, M., Schultz, C. J., Borner, G. H., Dupree, P., & Eisenhaber, F. (2003). Glycosylphosphatidylinositol lipid anchoring of plant proteins. Sensitive prediction from sequence- and genome-wide studies for Arabidopsis and rice. Plant Physiol, 133(4), 1691– 1701. 10.1104/pp.103.023580

Evans, R., O’Neill, M., Pritzel, A., Antropova, N., Senior, A., Green, T., Žídek, A., Bates, R., Blackwell, S., Yim, J., Ronneberger, O., Bodenstein, S., Zielinski, M., Bridgland, A., Potapenko, A., Cowie, A., Tunyasuvunakool, K., Jain, R., Clancy, E.,…Hassabis, D. (2021). Protein complex prediction with AlphaFold-Multimer. bioRxiv.

Ferreira, C., Silva, S., Faria-Oliveira, F., Pinho, E., Henriques, M., & Lucas, C. (2010). Candida albicans virulence and drug-resistance requires the O-acyltransferase Gup1p. BMC Microbiol, 10, 238. 10.1186/1471-2180-10-238

Gabaldon, T. (2021). Origin and Early Evolution of the Eukaryotic Cell. Annu Rev Microbiol, 75, 631– 647. 10.1146/annurev-micro-090817-062213

Gerber, L. D., Kodukula, K., & Udenfriend, S. (1992). Phosphatidylinositol glycan (PI-G) anchored membrane proteins. Amino acid requirements adjacent to the site of cleavage and PI-G attachment in the COOH-terminal signal peptide. J Biol Chem, 267(17), 12168–12173.

Hardy, R. Y., & Resh, M. D. (2012). Identification of N-terminal residues of Sonic Hedgehog important for palmitoylation by Hedgehog acyltransferase. J Biol Chem, 287(51), 42881–42889. 10.1074/jbc.M112.426833

Harrison, S. J., Mott, E. K., Parsley, K., Aspinall, S., Gray, J. C., & Cottage, A. (2006). A rapid and robust method of identifying transformed Arabidopsis thaliana seedlings following floral dip transformation. Plant Methods, 2, 19.

Hemsley, P. A., Hurst, C., Kaliyadasa, E., Lamb, R., Knight, M. R., De Cothi, E. A., Steele, J. F., & Knight, H. (2014). The MED16, MED14 and MED2 subunits of the Arabidopsis Mediator complex control recruitment of Mediator and RNA polymerase II to CBF-responsive cold-regulated genes. The Plant Cell.

Hurst, C. H., Turnbull, D., Xhelilaj, K., Myles, S., Pflughaupt, R. L., Kopischke, M., Davies, P., Jones, S., Robatzek, S., Zipfel, C., Gronnier, J., & Hemsley, P. A. (2023). S-acylation stabilizes ligand-induced receptor kinase complex formation during plant pattern-triggered immune signaling. Curr Biol, 33(8), 1588–1596 e1586. 10.1016/j.cub.2023.02.065

Jaquenoud, M., Pagac, M., Signorell, A., Benghezal, M., Jelk, J., Butikofer, P., & Conzelmann, A. (2008). The Gup1 homologue of Trypanosoma brucei is a GPI glycosylphosphatidylinositol remodelase. Mol Microbiol, 67(1), 202–212. 10.1111/j.1365-2958.2007.06043.x

Jiang, Y., Benz, T. L., & Long, S. B. (2021). Substrate and product complexes reveal mechanisms of Hedgehog acylation by HHAT. Science, 372(6547), 1215–1219. 10.1126/science.abg4998

Karimi, M., Inze, D., & Depicker, A. (2002). GATEWAY vectors for Agrobacterium-mediated plant transformation. Trends Plant Sci, 7(5), 193–195. S1360-1385(02)02251-3 [pii]

Kleinboelting, N., Huep, G., Kloetgen, A., Viehoever, P., & Weisshaar, B. (2012). GABI-Kat SimpleSearch: new features of the Arabidopsis thaliana T-DNA mutant database. Nucleic Acids Res, 40(Database issue), D1211–1215. 10.1093/nar/gkr1047

Lalanne, E., Honys, D., Johnson, A., Borner, G. H., Lilley, K. S., Dupree, P., Grossniklaus, U., & Twell, D. (2004). SETH1 and SETH2, two components of the glycosylphosphatidylinositol anchor biosynthetic pathway, are required for pollen germination and tube growth in Arabidopsis. Plant Cell, 16(1), 229–240. 10.1105/tpc.014407

Li, Y., Wang, X., Zhou, N. Y., & Ding, J. (2024). Yeast surface display technology: Mechanisms, applications, and perspectives. Biotechnol Adv, 76, 108422. 10.1016/j.biotechadv.2024.108422

Lucas, C., Ferreira, C., Cazzanelli, G., Franco-Duarte, R., & Tulha, J. (2016). Yeast Gup1(2) Proteins Are Homologues of the Hedgehog Morphogens Acyltransferases HHAT(L): Facts and Implications. J Dev Biol, 4(4). 10.3390/jdb4040033

Mirdita, M., Schütze, K., Moriwaki, Y., Heo, L., Ovchinnikov, S., & Steinegger, M. (2022). ColabFold: making protein folding accessible to all. Nature Methods, 19(6), 679–682. 10.1038/s41592-022-01488-1

Miura, G. I., Buglino, J., Alvarado, D., Lemmon, M. A., Resh, M. D., & Treisman, J. E. (2006). Palmitoylation of the EGFR ligand Spitz by Rasp increases Spitz activity by restricting its diffusion. Dev Cell, 10(2), 167–176. 10.1016/j.devcel.2005.11.017

Moore, R. M., Harrison, A. O., McAllister, S. M., Polson, S. W., & Wommack, K. E. (2020). Iroki: automatic customization and visualization of phylogenetic trees. PeerJ, 8, e8584. 10.7717/peerj.8584

Nakagawa, T., Suzuki, T., Murata, S., Nakamura, S., Hino, T., Maeo, K., Tabata, R., Kawai, T., Tanaka, K., Niwa, Y., Watanabe, Y., Nakamura, K., Kimura, T., & Ishiguro, S. (2007). Improved Gateway binary vectors: high-performance vectors for creation of fusion constructs in transgenic analysis of plants. Biosci Biotechnol Biochem, 71(8), 2095–2100. 10.1271/bbb.70216

Oxley, D., & Bacic, A. (1999). Structure of the glycosylphosphatidylinositol anchor of an arabinogalactan protein from Pyrus communis suspension-cultured cells. Proc Natl Acad Sci U S A, 96(25), 14246–14251. 10.1073/pnas.96.25.14246

Schmittgen, T. D., & Livak, K. J. (2008). Analyzing real-time PCR data by the comparative C(T) method. Nat Protoc, 3(6), 1101–1108.

Schonbrun, A. R., & Resh, M. D. (2022). Hedgehog acyltransferase catalyzes a random sequential reaction and utilizes multiple fatty acyl-CoA substrates. J Biol Chem, 298(10), 102422. 10.1016/j.jbc.2022.102422

Sessions, A., Burke, E., Presting, G., Aux, G., McElver, J., Patton, D., Dietrich, B., Ho, P., Bacwaden, J., Ko, C., Clarke, J. D., Cotton, D., Bullis, D., Snell, J., Miguel, T., Hutchison, D., Kimmerly, B., Mitzel, T., Katagiri, F.,…Goff, S. A. (2002). A high-throughput Arabidopsis reverse genetics system. Plant Cell, 14(12), 2985–2994. 10.1105/tpc.004630

Sievers, F., Wilm, A., Dineen, D., Gibson, T. J., Karplus, K., Li, W., Lopez, R., McWilliam, H., Remmert, M., Soding, J., Thompson, J. D., & Higgins, D. G. (2011). Fast, scalable generation of high-quality protein multiple sequence alignments using Clustal Omega. Mol Syst Biol, 7, 539. 10.1038/msb.2011.75

Sueda, S., Tsuruga, R., Hirakawa, T., & Fujii, S. (2025). Cell surface display of a protein based on a tail-anchored membrane protein. Biochem Biophys Res Commun, 761, 151738. 10.1016/j.bbrc.2025.151738

Svetek, J., Yadav, M. P., & Nothnagel, E. A. (1999). Presence of a glycosylphosphatidylinositol lipid anchor on rose arabinogalactan proteins. J Biol Chem, 274(21), 14724–14733. 10.1074/jbc.274.21.14724

Takada, R., Satomi, Y., Kurata, T., Ueno, N., Norioka, S., Kondoh, H., Takao, T., & Takada, S. (2006). Monounsaturated fatty acid modification of Wnt protein: its role in Wnt secretion. Dev Cell, 11(6), 791–801. 10.1016/j.devcel.2006.10.003

Takahashi, D., Kawamura, Y., & Uemura, M. (2016). Cold acclimation is accompanied by complex responses of glycosylphosphatidylinositol (GPI)-anchored proteins in Arabidopsis. J Exp Bot, 67(17), 5203–5215. 10.1093/jxb/erw279

Tashima, Y., Taguchi, R., Murata, C., Ashida, H., Kinoshita, T., & Maeda, Y. (2006). PGAP2 is essential for correct processing and stable expression of GPI-anchored proteins. Mol Biol Cell, 17(3), 1410–1420. 10.1091/mbc.e05-11-1005

Umemura, M., Fujita, M., Yoko, O. T., Fukamizu, A., & Jigami, Y. (2007). Saccharomyces cerevisiae CWH43 is involved in the remodeling of the lipid moiety of GPI anchors to ceramides. Mol Biol Cell, 18(11), 4304–4316. 10.1091/mbc.e07-05-0482

Van der Vaart, J. M., te Biesebeke, R., Chapman, J. W., Toschka, H. Y., Klis, F. M., & Verrips, C. T. (1997). Comparison of cell wall proteins of Saccharomyces cerevisiae as anchors for cell surface expression of heterologous proteins. Appl Environ Microbiol, 63(2), 615–620. 10.1128/aem.63.2.615-620.1997

Voinnet, O., Rivas, S., Mestre, P., & Baulcombe, D. (2003). An enhanced transient expression system in plants based on suppression of gene silencing by the p19 protein of tomato bushy stunt virus. Plant J, 33(5), 949–956. 1676 [pii]

Xu, Z., Gao, Y., Gao, C., Mei, J., Wang, S., Ma, J., Yang, H., Cao, S., Wang, Y., Zhang, F., Liu, X., Liu, Q., Zhou, Y., & Zhang, B. (2022). Glycosylphosphatidylinositol anchor lipid remodeling directs proteins to the plasma membrane and governs cell wall mechanics. Plant Cell, 34(12), 4778– 4794. 10.1093/plcell/koac257

Yan, P., Shen, W., Gao, X., Li, X., Zhou, P., & Duan, J. (2012). High-throughput construction of intron-containing hairpin RNA vectors for RNAi in plants. PLoS One, 7(5), e38186. 10.1371/journal.pone.0038186

Yang, J., Brown, M. S., Liang, G., Grishin, N. V., & Goldstein, J. L. (2008). Identification of the acyltransferase that octanoylates ghrelin, an appetite-stimulating peptide hormone. Cell, 132(3), 387–396. 10.1016/j.cell.2008.01.017

Yeats, T. H., Bacic, A., & Johnson, K. L. (2018). Plant glycosylphosphatidylinositol anchored proteins at the plasma membrane-cell wall nexus. J Integr Plant Biol, 60(8), 649–669. 10.1111/jipb.12659

